# Policy should change to improve invited speaker diversity and reflect trainee diversity

**DOI:** 10.1101/785717

**Authors:** Ada K. Hagan, Rebecca M. Pollet, Josie Libertucci

**Affiliations:** Department of Microbiology & Immunology, University of Michigan, Ann Arbor, Michigan; Farncombe Family Digestive Health Research Institute, Department of Medicine, McMaster University, Hamilton, Ontario, Canada

**Keywords:** inclusion, diversity, invited speakers, academia, graduate programs

## Abstract

The biomedical sciences have a problem retaining white women and underrepresented minorities in academia. Despite increases in the representation of these groups in faculty candidate pools, they are still underrepresented at the faculty level, particularly at the Full Professor level. The lack of diverse individuals at the Full Professor level contributes to the attrition of women and under-represented minorities, as it confirms unconscious biases. The presence of unconscious biases contribute to feelings of not belonging by trainees and are amplified by visual representation of who is presented as the “top scientist in their field”. Top scientists are not only defined by the attainment of Full Professorships, but also through invited seminar series. Invitations for faculty to present their research at other university departments is highly valued offer that provides an opportunity for collaborations and networking. However, if invited speakers do not represent the demographics of current trainees, these visual representations of successful scientists may contribute to decreased attitudes of self-identification as a scientist, ultimately resulting in trainees leaving the field or the academy. In this study, we compare invited-speaker demographics to the current trainee demographics in one microbiology and immunology department and find that trainees are not proportionally represented by speakers invited to the department. Our investigation prompted changes in policy for how invited speakers are selected in the future to invite a more diverse group of scientists. To facilitate this process, we developed a set of tips and a web-based resource that allows scientists, committees, and moderators to identify members of under-served groups. These resources can be easily adapted by other fields or sub-fields to promote inclusion and diversity at seminar series’, conferences, and colloquia.

## Background

Long-standing systemic bias, sexism, and racism have contributed to the under-representation of many racial and ethnic groups, as well as women, in science, technology, engineering, and math (STEM) fields (1–4). Specifically, within the field of biomedical research in the United States, the proportion of underrepresented minorities at the full professor level has remained consistently low at 4% (survey data taken from the NIH from 2001 to 2013), compared to the U.S. population, which is 32.3% (5, 6). Similar discrepancies exist for women in biomedical sciences as full professorships are currently held mostly by men (7, 8). As demographics of faculty within the biomedical sciences remains skewed towards Caucasian men, the demographics of trainees (graduate students and postdocs) are becoming more diverse (5).

Policy changes are needed to support inclusion of all individuals, particularly in the biomedical sciences(9). To increase retention of historically under-represented minorities (HURM), non-Caucasian/non-HURM (NCNH) individuals, and white women in biomedical fields, it is important for trainees to have visual representations of themselves as scientists. The importance of representation in retaining a diverse group of individuals in STEM fields is supported by social role theory (10). Individuals make inferences about characteristics that are needed to be successful in a given role by examining individuals that most occupy that role (10, 11). However, there is a lack of diverse scientific experts in academia so underrepresented minorities are not seeing adequate visual representations of themselves at the faculty level. Therefore, trainees who do not see representation of themselves in senior faculty positions, may decide that they do not possess the characteristics that are required to succeed.

Invited seminar series are common within biomedical departments across the United States (12). Usually, seminar series’ consist of faculty members selecting a scientist from another institution to visit their university and present their research, as well as meet with other faculty members and trainees. Named lectureships follow the same format but are decided by committee and are considered more prestigious because they are named in honor of prominent local scientists. These seminar series and lectureships provide an opportunity for trainees to be exposed to research outside of their department. Additionally, being an invited speaker provides the scientist with an opportunity to make future collaborations and build their own *curriculum vitae* (CV). Scientists invited to give seminars are widely regarded as successful and the top in their field. Thus, if trainees are constantly being exposed to “the top scientist in their field”, according to social role theory, it signals who is successful in that field. While some have examined this issue by studying and promoting the inclusion of more women speakers at conferences, how department speaker series compare to the trainee diversity of that department is unknown (13–15).

In this study, we examine and compare the proportion of HURM, NCNH, and women invited speakers to white men in the Department of Microbiology and Immunology at the University of Michigan. Additionally, we compare invited-speaker demographics to the current trainee demographics as a means to gauge if trainee demographics are being represented accordingly throughout the seminar series. Following our investigation, we proposed a policy change to the Department of Microbiology and Immunology in how invited speakers are selected as a means to promote inclusion in our department and reduce unconscious bias. In order to facilitate inviting a more diverse group of scientists, we developed a set of resources that allow scientists, within the fields of microbiology and immunology, to self-identify as having an under-represented or under-served identity including: HURM, non-Caucasian/non-HURM, or a white woman. These resources will promote inclusion and diversity by providing greater representation of all scientists and will provide hosts an opportunity to invite a more diverse group of scientists.

## Methods

Each academic year, each faculty member in the Department of Microbiology and Immunology at the University of Michigan has the opportunity to invite one speaker per year for a weekly seminar series. Some of these seminar slots are dedicated to named lectureships, which are decided by committee, and three trainee-invited speakers. We analyzed the demographics of invited speakers and faculty hosts for five academic years (Fall 2014 - Spring 2019), and compared them to the current trainees when the data were analyzed (Spring 2019). Each speaker was only counted once and those listed as departmental faculty members or as a “host” at any point could not also be considered “invited speakers”. The list of faculty hosts was used as a proxy for faculty demographics since as hosts, these faculty members are visible representatives of the department. The trainees were identified using departmental email lists that included masters students, doctoral students, and post-doctoral fellows.

This is a retrospective study, thus speakers were not asked for their identities at the time of visit. Instead we hand-coded proxy demographics using personal knowledge, photos, and CVs. The presenting gender of each individual was assigned using a binary system (man/woman). Due to the low number of individuals in the study, race/ethnicity demographics were split in three groups: Caucasian, Historically Under-represented Minority (HURM), and Non-Caucasian/Non-HURM (NCNH), each with a binary (yes/no) possibility. Caucasian was assigned using the current U.S. Census definition where those of Middle Eastern, European, and Russian descent are included. HURM individuals were restricted to those with African-American, Indigenous, Alaskan/Hawaiian Native, Latinx and/or Hispanic heritage. All others were placed into the NCNH group. We recognize that our proxy demographics are a limitation of the analysis and want to acknowledge that biological sex (male/female) is not always equivalent to the gender that an individual presents as (man/woman), which is also distinct from the gender(s) that an individual self-identifies as. We also want to acknowledge that there are many other identities that are not captured in this limited analysis.

Data were analyzed and figures generated in R Statistical Software, using relevant packages (16–28).

## Results

To understand the representation of women, we compared the proportion of women in each academic role. At the trainee level, more than half of students and postdoctoral fellows were women. That dropped to 46.77% of faculty hosts and 38.73% of the invited speakers (Fig. 1A). Of 27 lectureships over the five year period, 37.04% were awarded to women.

**Figure 1:**
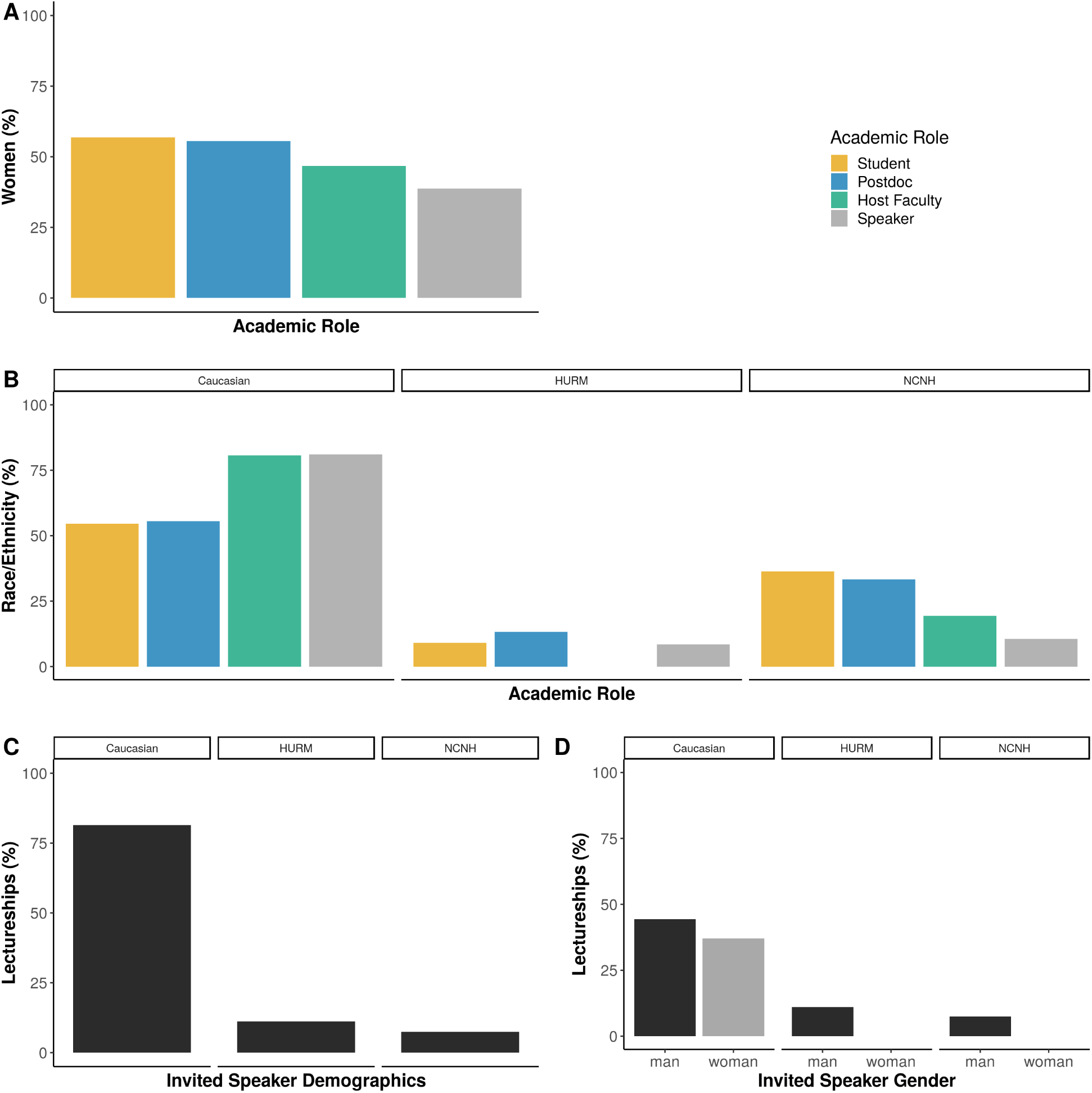
The demographics of invited speakers, hosting faculty, and trainees. A) The proportion of women in each academic role. B) The proportion of each academic role represented by individuals that are Caucasian (left), Historically Underrepresented Minorities (HURM, center) or Non-Caucasian/Non-HURM (NCNH, right). C-D)The percent of lectureships awarded to individuals that are C) Caucasian, HURM, or NCNH and D) Caucasian, HURM, or NCNH by gender.

Our analysis identified an over-representation of Caucasian individuals as hosting faculty and invited speakers (80% each), relative to the proportion of Caucasian trainees, which was 55% (Fig. 1B). We also observed declines in the representation of HURM and NCNH faculty and speakers relative to the trainees (Fig 1B). HURM trainees made up 11% of the department, on track with the 11% of U.S. microbiology and immunology doctorates awarded in 2017 (29). However, only 8.5% of invited speakers, and none of the hosting faculty, were HURM scientists. NCNH trainees were 34% of department students and postdocs (versus 22% of U.S. microbiology and immunology doctorates in 2017), but only 19% of hosting faculty and 10.5% of invited speakers (29).

The more prestigious invited speaker lectureships were also dominated by Caucasian scientists, who comprised 81.48% of those awarded (Fig. 1C). HURM and NCNH scientists were awarded 3 and 2 lectureships, respectively. Because the intersection of identities can compound biases and outcomes, we further examined the lectureships by gender and race/ethnicity status (30). Caucasian men and women accounted for 44.44% and 37.04% of the lectureships, respectively. Just 18.52% of lectureships were held by non-Caucasian men while none were held by non-Caucasian women (Fig. 1D).

## Discussion

This study found that the proportion of HURM and NCNH invited speakers were under-representative of the trainee populations for each group. Additionally, within the last 5 years, no HURM or NCNH woman was awarded a lectureship. This means that the department is not providing non-Caucasian trainees with adequate representation of successful scientists. Taking this into context of social role theory, by not adequately representing the diversity of all trainees, the department is not supporting an inclusive environment in terms of visual faculty representation. We also found that the proportion of women as faculty hosts and speakers in our study population is equivalent to global estimates that 40% of microbiologists are women, though women only represent about 30% of academic biomedical faculty (7, 31). Women are also over-represented as graduate students and postdoctoral fellows in this department. Overall, Caucasian scientists are over-represented as host faculty and invited speakers, compared to their presence as trainees, particularly when lectureships were considered.

Several papers have investigated the representation of women at scientific conferences, however, we have only identified one that focused on invited speakers at universities (12). In their study, Nittrouer et al, examined 3,652 talks at 50 U.S. institutions in 2013 - 2014 and found that women faculty are less likely to be invited speakers, despite similar acceptance rates (12). We have not been able to identify any publications examining scientific speaker diversity beyond gender. This seems to be the first, which is concerning since conclusions drawn from gender-based studies are often framed, and considered, to be applicable to other marginalized groups (e.g., HURM). This is a flawed assumption. While there is no doubt some overlap, each group remains marginalized due to a unique complex set of factors that cannot always be solved by gender-based solutions. U.S. institutions, such as the University of Michigan have a particular responsibility to the historically suppressed populations included in our definition of HURMs. We therefore implore U.S. institutions to apply this framing to their discussions and research.

Departments have different processes and criteria for selecting invited speakers, but it is a matter of pride to bring the best scientists possible. It may be that the definition of “best” poses a problem to under-represented and under-served groups (e.g., white women, HURM, and Asian) who are held to stricter competency standards and report having to work harder than white men to be perceived as legitimate scholars (32, 33). Some departments only invite tenured faculty, which severely limits the number of potential speakers who are white women or non-Caucasian. Yet, another scenario is that pre-tenure faculty members invite prestigious, tenured faculty in their field to network and secure letters for their own tenure package. The increased burden of white women and non-Caucasian scientists to prove competency decreases their likelihood to be considered for either tenure or as possible source of tenure letters.

Each underrepresented group in our cohort faces a complex set of barriers to achieving faculty status. For instance, the decision to invite a woman may also be negatively impacted by assumptions about competency and dedication. The dedication of women who have children to their work is perceived to be less than that of their colleagues, i.e., men who also have children (34–36). The perceived prioritization and commitments of women to family over work may cause faculty to doubt their acceptance of a speaking invitation, despite the prestigious nature of these invitations and evidence that men and women accept at similar rates (12, 37). As a result, the faculty member may invite a different colleague who they feel is more likely to agree (and is a man). Another large portion of our sample were the NCNH cohort, who are predominately Asian/Asian American individuals. Although Asian scientists are well-represented in the US scientific workforce, they face significant bias and barriers to inclusion in society and academia (38, 39). For instance, despite the higher employment rate of Asian scientists, they were not well-represented in the more prestigious lectureships.

While HURM and NCNH share some experiences, differences including varying rates of hiring and tenure promotion mean unique considerations are important for inclusion of each group (3). For instance, a major barrier to inclusion of HURM faculty at similar proportions to HURM trainees is the low transition rate of scientists from HURM backgrounds to faculty positions and the associated low proportion of HURM faculty (40). The proportion of HURM faculty at the Assistant and Associate Professor level is currently higher than at Full Professor so it will be difficult to increase speaker diversity if early-career researchers are not being considered (41). Increased performance expectations and patterns of exclusions are consistent themes in studies characterizing the HURM faculty experience (42, 43). Therefore, inclusion of HURM faculty in seminar series is likely essential to increasing the number of HURM Associate and Full Professors. Even when HURM speaker rates match the proportion of HURM faculty employment, HURM trainees will be represented at a significantly higher proportion. Inclusion of HURM faculty in these seminar series is just one aspect of larger institutional change that is needed (44).

## Instituting Policy Change

In an attempt to promote inclusion within the Department of Microbiology and Immunology at the University of Michigan, these data were presented to faculty members and the department chair (Dr. Mobley). Since trainee demographics were not represented by the seminar speaker demographics over the past 5 years, we proposed a policy change as to how seminar speakers were being invited. One suggestion was to switch from faculty-invited to lab-invited speakers in an attempt to allow trainees to choose a speaker that best represented themselves (Table 1).

**Table 1:** List of suggestions and resources to increase invited speaker diversity.

| Suggestion | Description | Resource |
| --- | --- | --- |
| Lab-invited speakers | Faculty members can request suggestions from trainees |  |
| Use a list | Many lists of scientists from under-represented and under-served groups are available | https://DiversifyMicrobiology.github.io/resources |
| Create a list | Use the GitHub template create a self-nomination list and resource for your field | https://github.com/diversifymicrobiology/DiversifyMicrobiology.github.io |
| Highlight the journey | Invite all speakers to spend a few moments describing their personal science journey |  |

The implicit biases that affect perceptions of marginalized groups are an issue, but we must acknowledge that it is not always possible to identify members of historically under-served communities. For instance, after data analysis, we learned that at least one speaker in our data set should have been categorized as a HURM instead of Caucasian, but it wasn’t readily apparent from their internet presence or CV. This limitation makes two important points: that perceived identity often plays a larger role than self-identification, and that we need better tools to identify members of marginalized groups. Another policy suggestion is for departments to invite their speakers to spend time discussing their personal journeys through science, in addition to their scientific stories (Table 1). This would enable those who wish, to discuss how their identity(ies) interacted with their careers. In addition to these suggestions for policy change, we have created resources that allow scientists to self-identify as under-served groups and thus provide host faculty with more diverse choices (Table 1).

## Building Diversify

Motivated by a lack of resources to identify scientists who are members of marginalized and/or historically under-served groups, and inspired by resources in other fields–DiversifyEEB and DiversifyChemistry–we created DiversifyMicrobiology and DiversifyImmunology (45–48). These resources are a tool for symposium organizers, award committees, search committees, and other scientists to identify individuals to diversify their pools. Additionally, we have built these as a template to be used by other fields and organizations that wish to create their own lists. Since these lists are compiled by self-nomination, we can ensure that only scientists comfortable revealing their marginalized identities are included.

The self-nomination form is a Google Form with entries logged in a private Google Sheet. This form is embedded within the website and can be linked to directly. The use of a Google Forms allows us to maintain this database at no cost and gives us the flexibility to add questions or change response options without disrupting previous responses. Entries are logged in a private spreadsheet so that entries can be screened before being added to the public database. This screening includes two steps: confirming that each person is listed in the database only once and that any submitted website is a personal, professional website. If both criteria are met, a new entry is added to the public database spreadsheet. If a person is already listed in the database, their information is updated to the most recent submission.

This public spreadsheet is embedded in the website and can be open separately as a locked (uneditable) Google Sheet, allowing the list to be easily searched. We have chosen to list individuals’ academic information first in the spreadsheet to encourage a focus on academic achievement rather than tokenization of marginalized identities. Currently the database lists individuals in order of self-nomination but future versions will be re-sorted based on name and/or academic field to varying the individuals who may receive more attention for simply being at the top of the list.

The website provides an interface to the Google forms and spreadsheets with template pages for viewing the list, adding a name to the list, and finding additional resources. Importantly, our website creation tool is hosted for free by GitHub, which provides a free website for each GitHub organization. Basic tools and skills required to set up a Diversify site include knowledge of, or experience with, the version control tool git, the web-tool GitHub, and a text editor. A tutorial in the DiversifyMicrobiology repository on GitHub provides links to these resources and instructions for adapting the tool to your own field (47).

## Conclusion

To increase the retention of white women, HURM and NCNH trainees in the biomedical sciences, they must also be represented as experts. However, the invited speaker diversity at one department does not represent the diversity of trainees. To facilitate the identification and recruitment of individuals in these historically under-served groups, we have built a tool to create self-nominated, field-specific lists.

## Acknowledgements

We thank Drs. Beth Moore and Harry Mobley and the Department of Microbiology & Immunology, University of Michigan for their input and financial support that enabled publication of our manuscript. We thank Bonnie Krey and former speaker series coordinators Drs. Nicole Koropatkin and Kathy Spindler for access to invited speaker data. We would also like to acknowledge and thank Nick Lesniak and Dr. Ariangela Kozick for their comments and suggestions.

## Author Contributions

A.K.H. collected the data, assigned demographics, analyzed the data, created the website, and wrote the methods and results. R.M.P. created the Google lists, forms, and website content and the description of their maintenance. J.L. wrote the introduction and provided conceptual advice. A.K.H. and J.L facilitated the policy change to the UM Department of Microbiology and Immunology. All authors contributed to preparing the final manuscript.

## Code and data availability

The anonymized data, code for all analysis steps, and an Rmarkdown version of this manuscript is available at https://github.com/akhagan/Hagan_SpeakerDiversity_JMBE_2019. Template and complete instructions for generating a field-specific Diversity website are available at https://github.com/diversifymicrobiology/DiversifyMicrobiology.github.io/.

